# Population genomics of *Marchantia polymorpha* subspecies *ruderalis* reveals evidence of climate adaptation

**DOI:** 10.1101/2024.11.19.624281

**Authors:** Shuangyang Wu, Katharina Jandrasits, Kelly Swarts, Johannes Roetzer, Svetlana Akimcheva, Masaki Shimamura, Tetsuya Hisanaga, Frédéric Berger, Liam Dolan

## Abstract

Sexual reproduction results in the development of haploid and diploid cell states during the life cycle. In bryophytes the dominant multicellular haploid phase produces motile sperm that swim through water to the egg to effect fertilization from which a relatively small diploid phase develops. In angiosperms, the reduced multicellular haploid phase produces non-motile sperm that is delivered to the egg through a pollen tube to effect fertilization from which the dominant diploid phase develops. These different life cycle characteristics are likely to impact the distribution of genetic variation among populations. However, little is known about the distribution of genetic variation among populations of bryophytes. To understand how genetic variation is distributed among populations of a bryophyte and to establish the foundation for population genetics research in bryophytes, we described the genetic diversity of collections of *Marchantia polymorpha* subspecies *ruderalis*, a cosmopolitan ruderal liverwort. We explored genetic diversity of this species using 78 genetically unique (non-clonal) accessions from a total of 209 collected from 37 sites in Europe and Japan. There was no detectable population structure among European populations but significant genetic differentiation between Japanese and European populations. By associating genetic variation across the genome with global climate data, we identified summer temperature and precipitation as climate factors influencing the frequency of adaptative alleles. We speculate that the requirement for water through which motile sperm swim imposes a constraint on the life cycle to which the plant genetically adapts.

## INTRODUCTION

Sexual reproduction results in the development of haploid and diploid cell states during the life cycle. Land plants develop multicellular stages in both the haploid and diploid phases of the life cycle. The haploid phase of the life cycle is dominant over the diploid phase among plants in one of the two monophyletic groups of land plants, the bryophytes. The diploid phase is dominant over the haploid phase in the other monophyletic group, the vascular plants which includes the angiosperms. The relative contributions of the haploid and diploid phases to the life cycle are likely to have consequences on the pattern of genetic diversity in bryophytes and vascular plants [1].

A great deal is known about the distribution of genetic diversity and adaptation among populations of angiosperms, a monophyletic group within the vascular plants (see for example [2–4]). The angiosperm egg is fertilized by a non-motile sperm delivered to the female by a pollen tube [5]. By contrast the bryophyte egg is fertilized by a motile sperm which requires liquid water to effect fertilization [6]. However, little is known about the distribution of genetic diversity among populations in bryophytes – plants in which the dominant stage in the life cycle is haploid and the diploid is produced by a flagellated sperm swimming to the egg to effect fertilization.

*Marchantia polymorpha* subspecies *ruderalis* is a liverwort with a haploid dominant life cycle, typical of a bryophyte [7]. It is an experimental model genetic system with a well annotated genome of approximately 218 Mb [8]. It is globally distributed in the Northern hemisphere and grows in a diversity of climates, often in ruderal communities. There is likely considerable variation in the ways in which these plants adapt to climate and other environmental factors. It is therefore a system in which we can investigate how genetic variation is distributed among populations both locally and globally.

*M. polymorpha* subspecies *ruderalis* is a ruderal and grows in disturbed habitats with open ground [9–11]. This includes disturbed open ground caused by human activity in urban areas, farms and transport infrastructure. It can also colonize open ground that forms after natural disturbance such as the clearings that form after forest or peatland fires [12]. Colonizing populations can expand rapidly through asexual reproduction or a combination of sexual and asexual reproduction [13]. If populations are derived from a single individual census number can increase through asexual reproduction and all individuals in the populations would be genetically identical (clones). Alternatively, populations may be derived from at least one male and one female, and the population would increase through both asexual and sexual reproduction. The resulting populations would include individuals with different genotypes produced through sexual reproduction, and each of these genetically unique individuals could also reproduce asexually producing sub-populations of clones.

The patterns of genetic variation and adaptation may be different in plants with a dominant haploid phase like *M. polymorpha* than in plants with a dominant diploid phase like angiosperms. It has been suggested that purging selection may be stronger in the haploid dominant life cycle than in the diploid dominant life cycle [1]. Furthermore, the diaspores – dispersal units – are different between angiosperms and bryophytes and may impact the distribution of genetic diversity. In *M. polymorpha*, there are two types of diaspores, spores and gemmae. Spores are produced by meiosis. They are approximately 10 µm in diameter and produced in their millions on individual plants [6, 14]. Their small size allows them to be carried over long distances by air and water currents enabling long distance gene flow. Gemmae are larger – approximately 500 µm in diameter and disc shaped – are clonal propagules derive from single cells in the epidermis of an adult plant [6]. While gemmae could be transported by water they would be too heavy to be carried on air currents. The production of diaspores that could disperse both locally (gemmae) and over very long distances (spores) might be expected impact genetic differentiation of populations of *M. polymorpha* on a landscape scale. Genome sequences are available for some *M. polymorpha* subspecies ruderalis accessions indicating that there is genetic variation among populations [15, 16]. However, it is unclear how genetic variation is distributed among these populations and in the landscape.

To discover the patterns of natural genetic variation in a species with a dominant haploid life cycle phase, we described the genetic diversity and population structure in *Marchantia polymorpha* subspecies *ruderalis*. We collected multiple individuals from unique populations locally in Europe and in Japan to ascertain the amount of genetic diversity. We supplemented this with single individuals from unique populations to get a global picture of patterns of genetic diversity. Sequencing 209 accessions identified 78 unique individuals; the remainder were genetically identical clones. The sampling of multiple individuals from local patches indicated that there is considerable genetic variation in sexually reproducing local populations. While analysis of sequence diversity of the entire collection indicated that the European and Japanese accessions are genetically differentiated, we discovered there is no population structure in Europe.

## RESULTS

### 1. Geographical distribution of 209 *Marchantia polymorpha* subsp. *ruderalis* accessions

To capture the genetic diversity of natural populations of *Marchantia polymorpha* spp. *ruderalis* accessions, we collected a total of 209 from 37 geographical locations (Figure 1A, Table S1), spanning in North America, Japan, and Europe. We collected both individuals from single sites, and multiple individuals from single sites (populations). We sequenced the genomes of all 209 accessions to an average coverage of 23.45X with a 94.08 % mapping rate for the nuclear genomes (Table S2). A total of 3,775,179 autosome SNPs were identified in the sample cohort with joint genotyping followed by stringent filtering[17]. Multiple individuals were collected from the same population in twelve sites in Europe (Table S1). Since *M. polymorpha* reproduces both vegetatively (through the production of propagules called gemmae) and sexually (though the production of spores produced by meiosis), we hypothesized that the curated collection of 209 individuals would include some clonally related individuals produced by vegetative reproduction and therefore genetically identical to each other.

**Figure 1.**
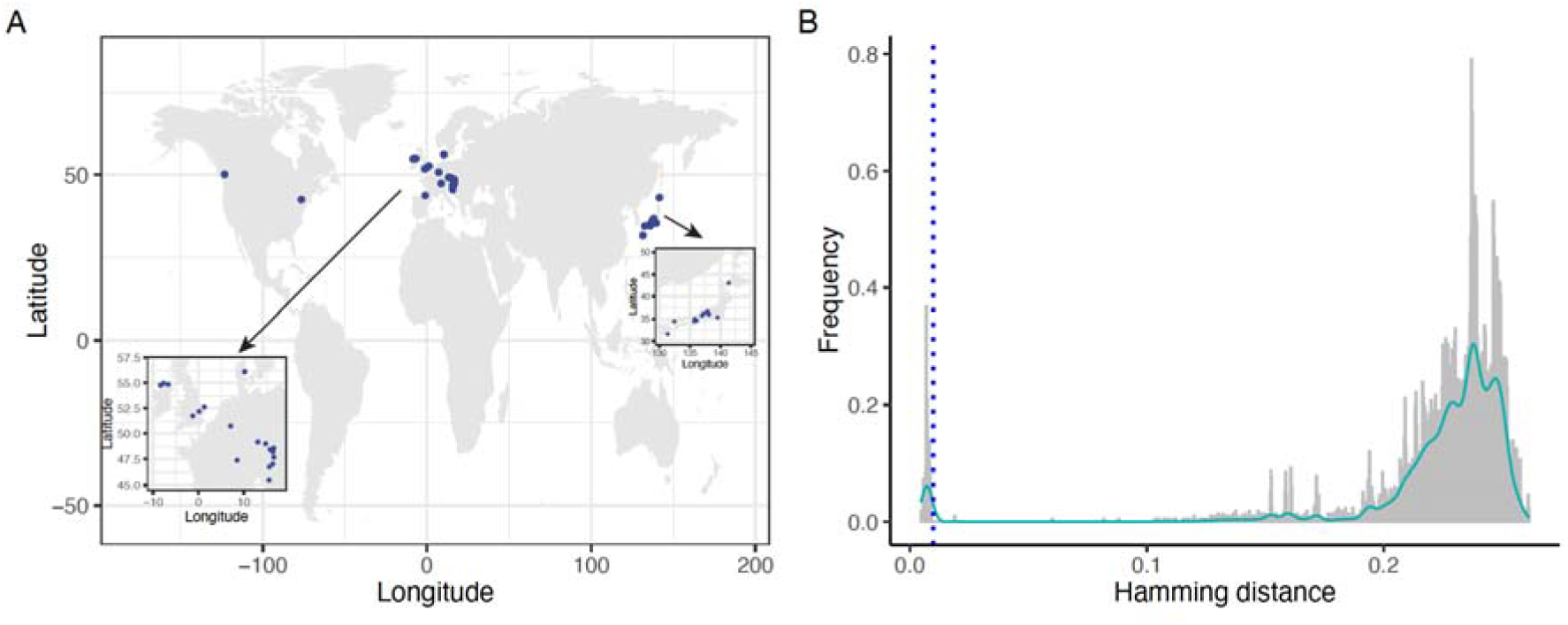
Geographical Distribution of 209 *M. polymorpha* subsp. *ruderalis* accessions and pairwise genome distance. A) Geographical distribution of 209 *M. polymorpha* subsp*. ruderalis* accessions. The blue dots represent the 37 collection sites. The small box on the left shows a zoomed-in view of the European sites, while the small box on the right shows a zoomed-in view of the Japanese sites. B) Pairwise genome distance distribution of 209 accessions, blue dashed line is the cutoff of clonal populations.

To identify clonal individuals, we calculated the Hamming distance as the pairwise genome distance among the 209 accessions, using all autosomal SNPs with Plink. (Table S3). We identified two distinct peaks in the GD distribution (Figure 1B), with peak A corresponding to a GD of less than 0.01, and peak B corresponding to a GD of 0.24. We plotted GD distribution in each population, and a similar pattern was observed (Figure S1). We conclude that pairs of accessions with GD values less than 0.01 (peak A) were clones. Using this criterion, amongst the twelve European populations studied two populations, Sopron (HUN) and MaG (AUT – with GD values at 0.008 and 0.0091, respectively – were entirely clonal; each population comprised a single genotype respectively (Figure S1). The other ten populations represented in peak B with higher GD (0.24) included individuals from each of the other sampled populations or individuals where a single plant was collected from a population. We conclude that most populations studied in Europe comprised individuals that were either genetically unique or clones, suggesting that some individuals in these populations are derived from sexual reproduction and others from vegetative reproduction (GD<=0.01). Isolated plants with no information regarding the diversity of the population but originating from a larger number of locations showed overall a GD> 0.1 and thus their genome was sufficiently distinct from each other to be considered as unique in our collection. Taken together, this demonstrates that some populations are entirely clonal (Figure S1), where each individual is derived from a single ancestor through vegetative reproduction. Other populations comprise both genetically unique individuals derived from sexual reproduction and clones produced by vegetative reproduction. In summary, we identified 78 genetically unique accessions originating from 36 diverse geographical locations (Table S4). This refined dataset now serves as a robust foundation for elucidating the genetic diversity present within Marchantia across varied global landscapes.

### 2. Population structure of 78 accessions reveals that Japanese and European populations are genetically distinct

Population structure is the differences in the allele frequencies between subpopulations of a population [18]. To explore the broad-scale population structure among accessions, we performed a principal component analysis (PCA) of the 3,479,055 pure autosome SNPs that were identified from the 78 samples through joint genotyping. Three genetically distinct clusters were identified: Japanese (Left bottom), Cambridge (Right top), and a mixed origin cluster (Right bottom) (Figure 2A). According to PC1, the structure can be divided into two groups: Japanese and European (including the 2 isolates from US and Canada). Further examination of the European cluster revealed samples that grouped together independent of country of origin (location) (Figure 2B), suggesting that there is no population structure among European accessions. To validate the PCA results, we performed Admixture analyses. Cross-validation (CV) error analysis from Admixture indicated that the optimal grouping was achieved at K=2 (Figure 2C). Two distinct groups were identified based on the genetic profile of the complete collection of accessions and their geographical origins (Japan versus Europe and north American continent) (Figure 2D). When we focused the admixture analysis on the European populations, we found mixed genetic ancestries of European populations from K3 to K9 (Figure S2A), indicating a lack of distinct population structure within European populations. In two European populations we sampled, Schubert (7 individuals) and Ardara (9 individuals), we found an equal number of ancestries compared to the broader European populations (From K7 to K9), which aligns with the hypothesis of high genetic diversity at the local level (Figure S2B). The absence of population structure among the European populations, as indicated by the admixture analysis, also aligns with the findings from the PCA analysis (Figure 2A, Figure 2B). In summary, these data demonstrate that the Japanese and European populations are genetically different. However, we did not detect any evidence for population structure among the European populations.

**Figure 2.**
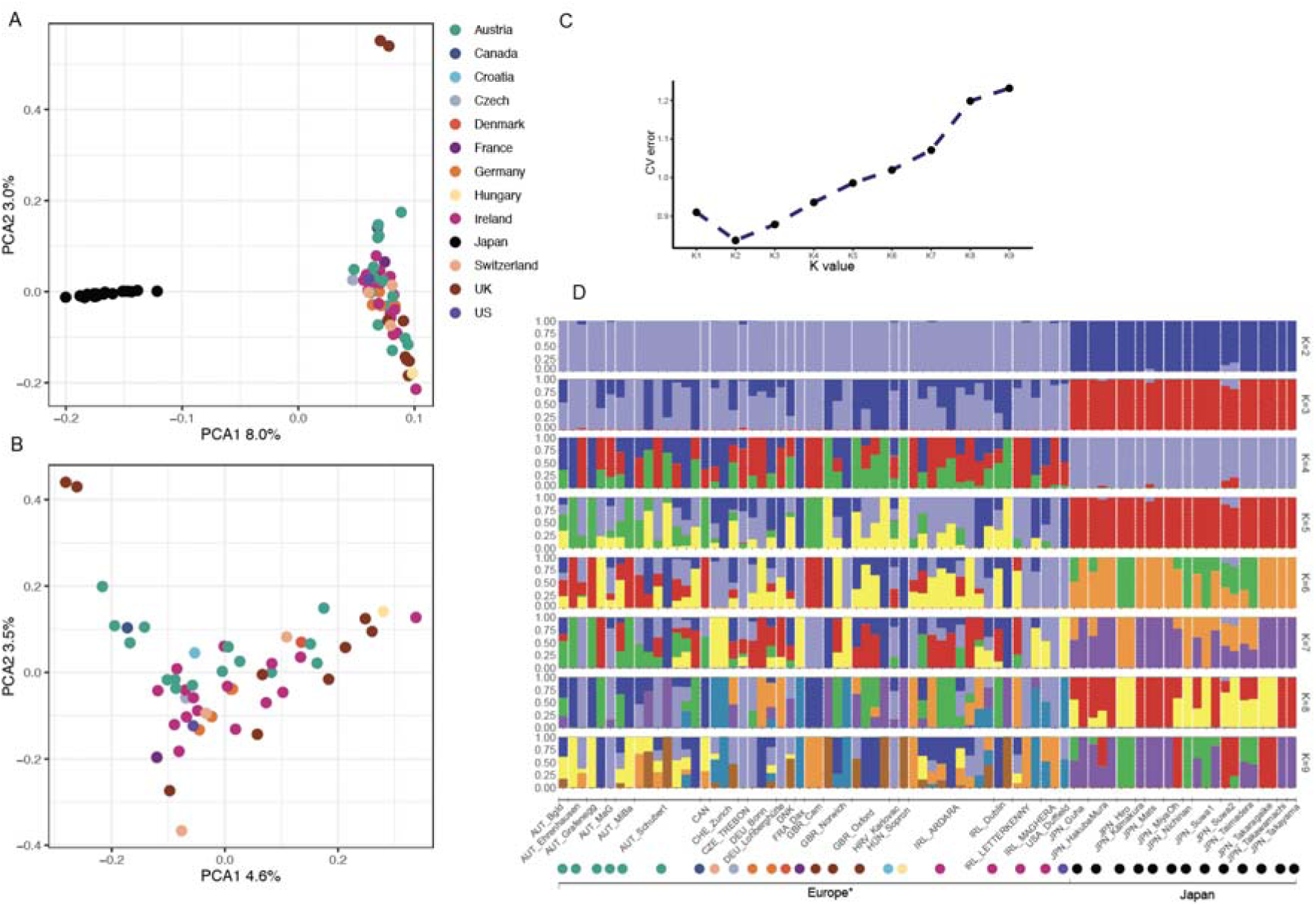
Genetic relationship of collected accessions. A) Principal component analysis (PCA) of all SNPs revealed 3 groups including Japanese (Left bottom), Cambridge (Right top), and a mixed origin cluster (Right bottom). B) PCA of European SNPs revealed no population structure within European populations. C) Cross-validation (CV) error analysis of the admixture result, demonstrating the optimal number of groups. The optimal K is 2. D) Genetic profile of the 78 accessions from K2 to K9. Dots below are color-coded by country, consistent with the color scheme in panel A. *two accessions from Canada and USA.

### 3. There is no substantial genetic differentiation among the European *Marchantia polymorpha subsp.* ruderalis populations

To test if there was genetic diversity difference between the European populations, we measured the nucleotide diversity (Pi, nucleotide difference within populations) for 7 populations, where 3 or more individuals were collected. Single representatives of clones were included in the analysis. Autosome SNPs were identified and the nucleotide diversity for each of these populations was close to 0.0036; Bonn_DEU (n=3), Ardara_IRL (n=9), Letterkenny_IRL (n=3), Oxford_GBR (n=4), Norwich_GBR (n=3), Schubert_AUT (n=7), and Zurich_CHE (n=3) (Figure 3A, Table S5). After assessing nucleotide diversity between these populations (Dxy, nucleotide diversity between populations), we found that the Dxy values among the seven populations were approximately 0.0042 (Figure 3B), which is close to the Pi value within each population. This indicates that there is low divergence in genetic diversity between these European populations.

**Figure 3.**
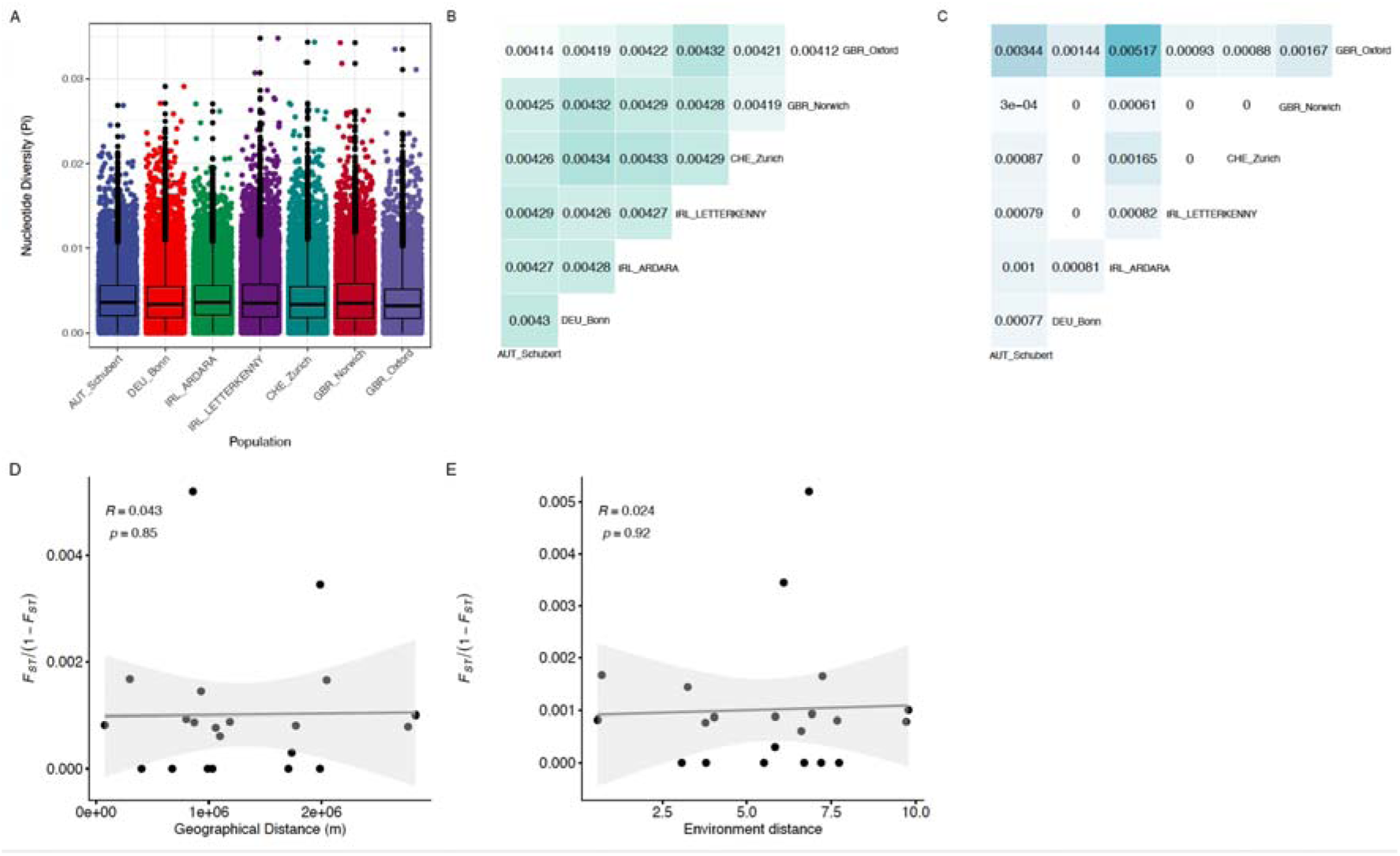
Genetic diversity and differentiation among the seven European populations. A) Genetic diversity among population groups was assessed by calculating nucleotide diversity in 20 kb sliding windows across the autosomes, with the median value indicated by the horizontal line. B) Pairwise median Dxy values between populations are represented by color-coded values, reflecting the genetic diversity between them. C) Pairwise mean Fst values between populations are depicted using a color scale, illustrating the extent of genetic differentiation between population pairs. D) Isolation-by-Distance analyses conducted across the seven European populations. The shadow of the linear regression line represents the 95% confidence interval. E) Isolation-by-Environment analyses performed within the seven European populations. The shadow of the linear regression line signifies the 95% confidence interval.

To test if genetic differentiation exists between these populations across Europe, a fixation index (Fst) analysis was performed to measure the degree of differentiation between pairs of the 7 populations, resulting in 21 Fst values from all possible pairwise comparisons. The pairwise Fst results (Figure 3C) indicated that the highest Fst value observed was 0.00517 indicating that there was no significant differentiation between the European populations. In *Arabidopsis thaliana* and other angiosperm species, genetic differentiation tends to increase with geographic distance (I.e Fst between North Sweden and South Sweden is ∼0.2[2]. However, our findings do not support this hypothesis. To independently test the hypothesis that genetic distance does not increase with geographic separation, an Isolation By Distance (IBD) analysis was performed between the seven populations. The results revealed no significant positive correlation of isolation by distance between the European populations, as determined by the Mantel test (Figure 3D). Since the accessions were collected from different parts of Europe, we carried out an Isolation By Environment (IBE) analysis. This analysis assesses the correlation between genetic distance and environmental distance. Again, there was no positive correlation between genetic distance and environmental distance, indicating the absence of significant isolation by environment (Figure 3E). Together, these data indicate that there is no substantial genetic differentiation among the European populations sampled.

### 4. There is substantial genetic differentiation between the European and Japanese *Marchantia polymorpha subsp. ruderalis* populations

Using the dataset from 54 European and 22 Japanese accessions we found that these two groups were genetically distinct from each other. Genetic diversity (Pi) was higher among the European accessions than among the Japanese accessions across 8 autosomes (Figure 4A, Figure S3 A). The Pi calculated from the autosomes of the European accessions was 0.00375, and 0.00246 in Japanese accessions (Figure S3 B). The difference in Pi values between the European and Japanese populations was not due to the larger sample size of the European group. This was further confirmed using bootstrap methods, with equal sample sizes ranging from 10 to 21 individuals from both Japan and Europe. Having shown that Pi is different between European and Japanese accessions, we then tested if there is genetic diversity divergence between these two populations using Dxy. Dxy between the two populations was 0.00403, which is far larger than the Pi value of Japanese accessions. This indicates that there is genetic differentiation between the two groups of accessions. To test genetic differentiation between European and Japanese groups of accessions, we calculated the Fst value. The genome-wide Fst between Japanese and European populations was 0.137. This is substantially higher than the within-region population differentiation values reported (with a mean of 0.0081for Europe), and quantifies the differentiation revealed by the in PCA and Admixture analysis (Figure 4B, Figure S3 C). This indicates that there is substantial genetic differentiation between the two – European and Japanese – groups of accessions.

**Figure 4.**
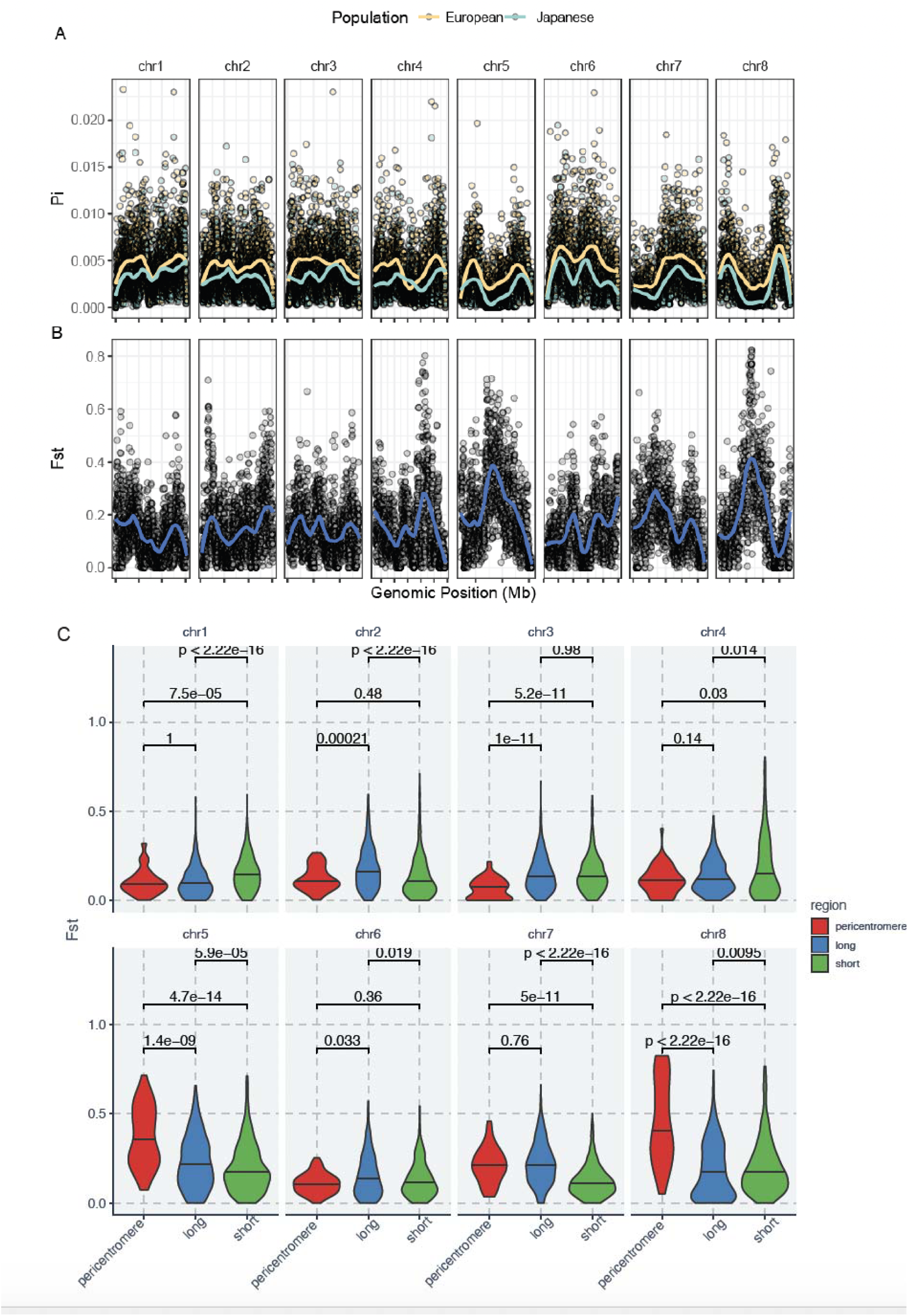
Chromosomal patterns of Marchantia genetic diversity and differentiation. Genetic diversity and differentiation were assessed by employing a sliding window approach with a window size of 20 kb. Each dot in the panels correspond to the value for each window. A) Genome-wide nucleotide diversity (Pi) for European (yellow) and Japanese (green) populations. Each dot represents the Pi value for a sliding window of 20 kb. B) Genome-wide Fst distribution between European and Japanese populations. Solid lines depict LOESS smoothing functions that have been fitted to the data, providing the representation of trends and patterns. C) The Fst distribution was analyzed for three segments of each autosome, the pericentromere, the long arm, and the short arm of chromosomes.

To identify which chromosome contributes to the genetic differentiation between European and Japanese groups, the Fst values for each chromosome were calculated. Chromosomes 5 and 8 contribute to a higher degree of the genetic differentiation than other autosomes (Figure 4B, Figure S3 D). This higher Fst values along chromosomes 5 and 8 suggested that the pericentromeres enriched in non-protein coding sequences contributed more to the genetic differentiation than the arms (Figure 4A) and chromosomes 5 and 8 had distinct centromeric haplotypes. To verify this, we partitioned each autosome into pericentromeres, long arm, and short arm. The analysis, as depicted in Figure 4C, demonstrated significantly higher differentiation in the pericentromeres of chromosomes 5 and 8 compared to the arm regions of these chromosomes. These data are consistent with the hypothesis that sequences in the vicinity of the centromeres of chromosomes 5 and 8 contribute disproportionately to differentiation between the Japanese and European accessions. In conclusion, Japanese and European populations are genetically distinct, and sequence variation in the vicinity of the centromeres of chromosome 5 and chromosome 8 contribute to the high levels of genetic differentiation (higher Fst) (Table S6).

### 5. Linkage disequilibrium and population demography of Japanese and European populations

Linkage disequilibrium (LD) refers to the tendency of genetic variants located near each other on a chromosome to be inherited together more frequently than those that are farther apart. LD is an essential measure to test if there are sufficient genetic markers in the populations for association mapping. To measure LD, we first measured LD decay using allele frequency correlation (r^2^) between linked SNPs markers. LD was then measured as the pair-wise SNP distance where *r*^2^ is half the maximum value[19, 20]. LD decay rate was higher in European groups than in Japanese groups (Figure 5A). From the decay plot (Figure 5A), LD of European population was estimated to be 1.6 kb, r^2^=0.1694) while the LD of the Japanese population was estimated to be 4.3 kb, r^2^=0.3262). LD for the entire collection of 78 individuals was estimated to be 1.5 kb. This estimation of LD is lower (1.5 Kb) than in Arabidopsis (LD 10 Kb) and rice (LD 150 Kb) [3, 21]. The minimum number of genetic markers required for genome wide association (GWA) can be estimated by dividing the genome size by the LD. The autosome size of Marchantia is ∼218 Mb. Therefore, the minimum number of SNPs is 218Mb/1.5 Kb = ∼ 145334 (n). Since we identified 3,479,055 SNPs in the collection, we conclude that there are sufficient genetic markers in Marchantia natural populations to perform GWA analysis.

**Figure 5.**
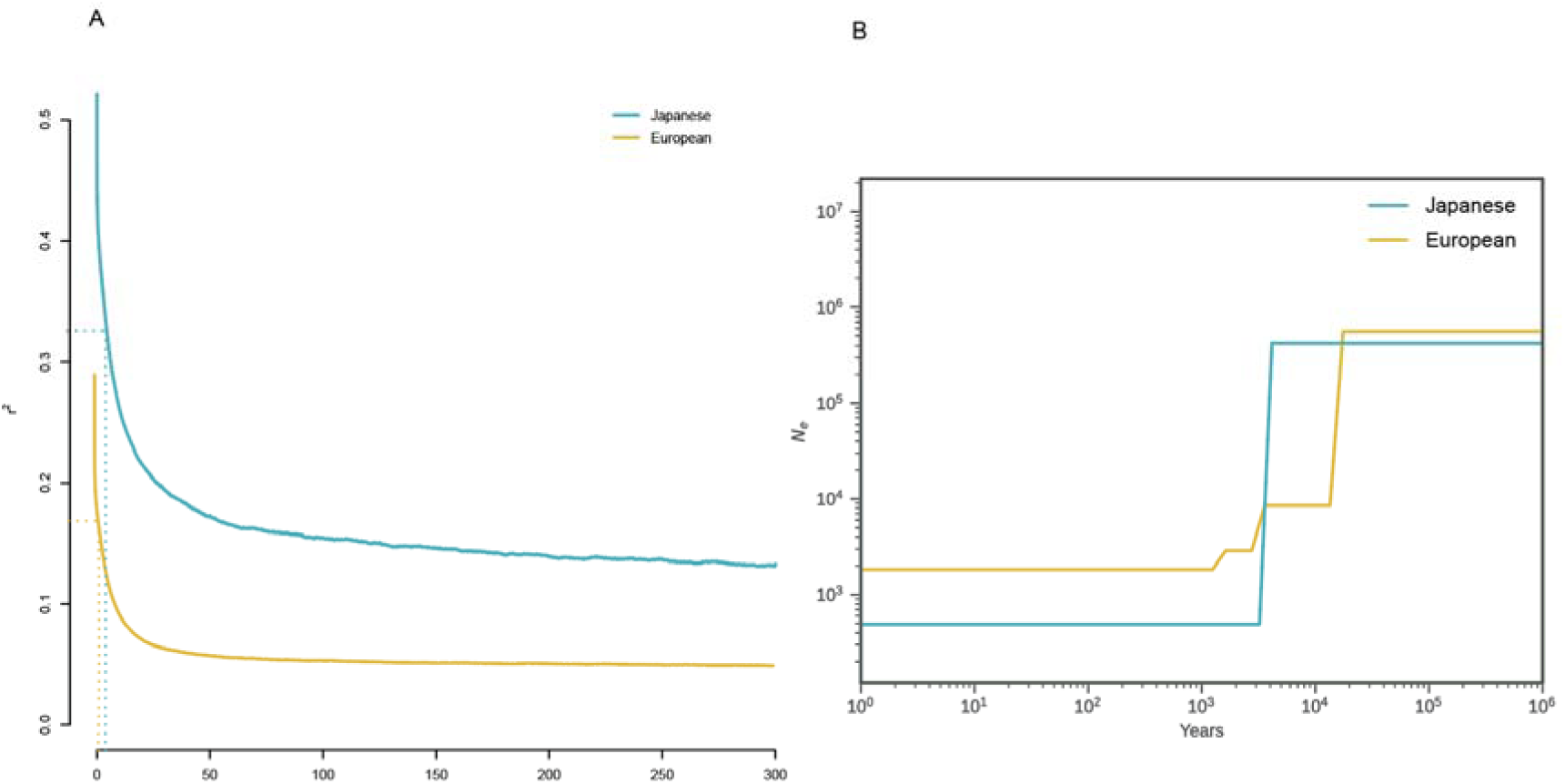
The LD decay and population demographic history of each population. A) LD decay in Japanese and European populations. Blue Solid line is LD decays plot for Japanese groups and yellow solid line is LD decay plot for the European groups. The LD decay rate of the European group is greater than in the Japanese group. The blue dashed line marks the X-intercept at half maximal r^2^ for the Japanese group (LD). The yellow dashed line marks the X-intercept at half maximal r^2^ for the European group (LD). B) Effective population sizes (Ne) distribution in Japanese groups (solid blue line) and European groups (solid yellow line).

Relatively low LD indicates larger effective population sizes, while higher LD in a population indicates a reduced effective population size. The lower LD in European populations compared with Japanese populations (Figure 5A) indicated that the effective population size is smaller in Japan than in Europe. To further test this idea, we conducted population demographic analyses to characterize the effective population size of European and Japanese populations (Figure 5B). Our results revealed that the current Marchantia effective population sizes (Ne) are significantly smaller than ancestral populations (10^6^ years ago). We estimate that the European Ne declined dramatically approximately 13,000 years ago, while the Ne declined approximately 5,000 years ago in Japan. Furthermore, the current effective population size of European (Ne = ∼3000) is larger than the current Japanese effective population size (Ne = ∼700). This result suggests that there may be more recombination occurring in the European population compared to the Japanese population at present.

### 6. Summer temperature and precipitation may be the most important climate factors affecting the frequency of adaptive alleles in European and Japanese accessions

To test for evidence of adaptation for ecological factors, we searched for association of markers with local climate data. WorldClim2 is a global climate database that provides high-resolution data for various environmental variables across the world, including 11 temperature factors and 8 precipitation factors [22]. We use GWAS in combination with global climate data to test if there is genetic adaptation to specific climate factors. Integrating allele frequencies and climatic data using the Gradient Forest (GF) method [23], we sought to identify environmental variables correlated with genetic variation and assessed how allele frequencies changed along environmental gradients. A total of 2,191 climate-associated genetic variants (SNPs) were identified using Latent Factor Mixed Models (LFMM)[24]. We calculated the allele frequencies of 2,191 SNPs in our European and Japanese groups. Combined with the 19 climate factors, we performed GF analysis with the 2,191 allele frequencies. Following GF ranking, five variables emerged as the most significant climate factors: Temperature Seasonality (BIO4), Precipitation of Warmest Quarter (BIO18), Maximum Temperature of Warmest Month (BIO5), Mean Temperature of Warmest Quarter (BIO10), and Mean Temperature of Wettest Quarter (BIO8) (Figure 6A). Maximum Temperature of Warmest Month (BIO5), Mean Temperature of the Warmest Quarter (BIO10), and Mean Temperature of the Wettest Quarter (BIO8) were equally ranked in importance. These data indicate that warm temperature and precipitation are the key climate factors influencing allele frequencies.

**Figure 6.**
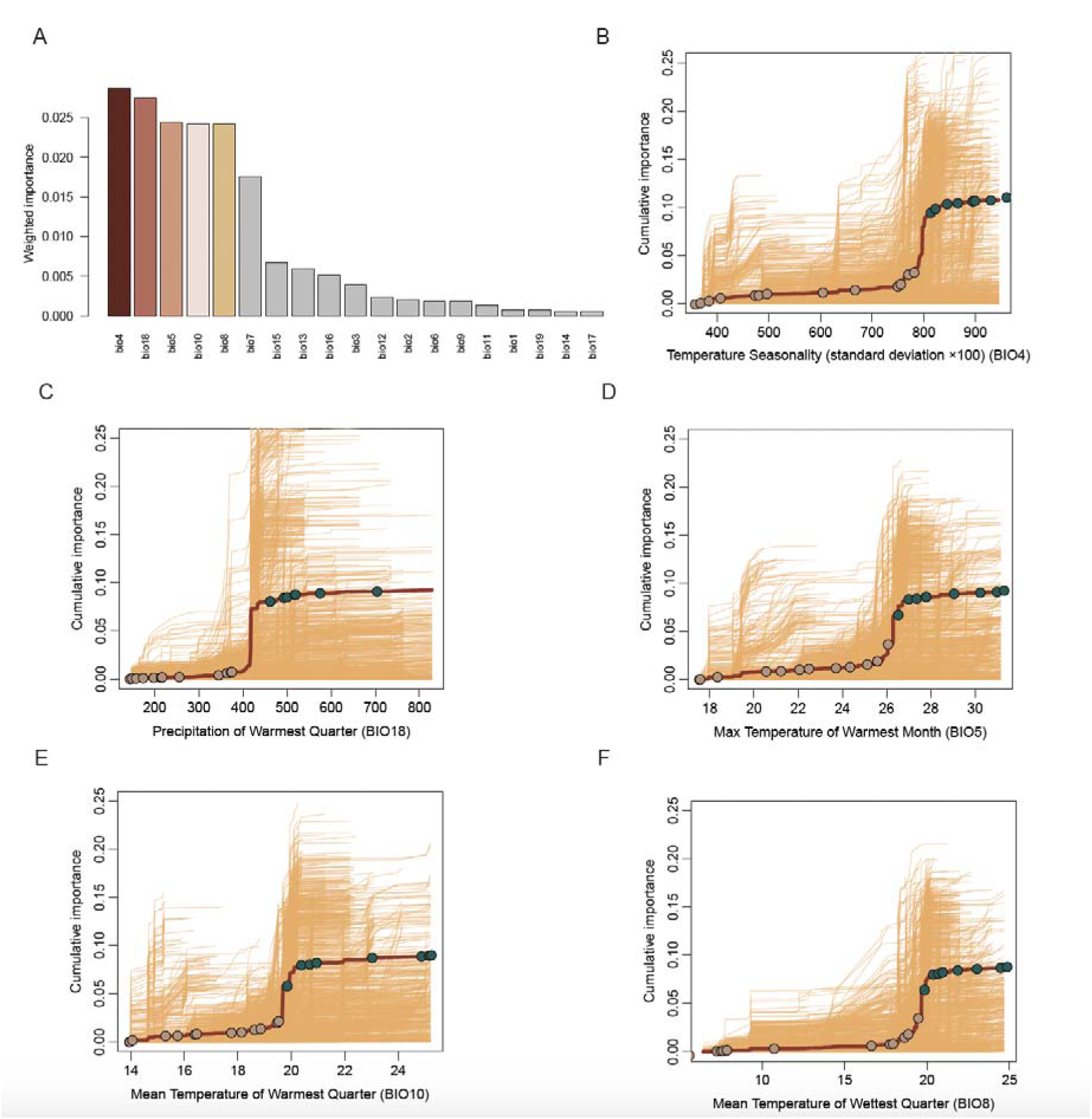
Gradient Forest and turnover function in climate factors. A) Ranked importance of 19 climate factors based on gradient forest analysis using 2191 candidate SNPs. The top five bioclimatic factors in the color gradient demonstrate the strongest inferred correlation. B - F) Allelic turnover functions in relation to bioclimatic factors BIO4 (B), BIO18 (C), BIO5 (D), BIO10 (E), and BIO8 (F) on the x-axis. The y-axis represents cumulated importance of allele frequencies, indicating the significance of SNPs in the GF models and reflecting the total turnover in allele frequency across the seasonality gradient. Thin yellow lines depict cumulative turnover for adaptive SNPs. The solid red line indicates turnover across all candidate SNPs, with circles along the line representing geographical groups organized based on different bioclimatic factors. Colors denote their inclusion in either the Japanese (green) or the European geographical groups (brown).

According to GF analysis, 4 temperature factors and 1 precipitation factor are the most influential climate variables. These 5 climates variables are likely relevant to the summer period. To test this hypothesis, we performed pairwise Spearman correlation analysis among the 19 climate factors. The pairwise Spearman correlation coefficients of BIO4, BIO5, BIO10, and BIO8 exceeded 0.85, indicating a significant pairwise positive correlation, suggesting these 4 temperature factors are associated with warm season temperature (Figure S4). Furthermore, these four factors were significantly negatively correlated with BIO6 (Minimum Temperature of Coldest Month) and BIO11 (Mean Temperature of Coldest Quarter), consistent with the positive association with Warm Season Temperatures. Furthermore, all four temperature variables (BIO4, BIO5, BIO10 and BIO8) were positively correlated (**) with BIO18, suggesting synergistic effects of Precipitation of Warmest Quarter with the warm season temperature. We conclude that all 5 factors highlight the same climate factor – summer temperature and precipitation – may be the most influential climate factor affecting the frequency of adaptation alleles.

To test if geographical groups were adapted to specific local environments, we integrated 2,191 SNPs allele frequencies and the 5 most important climate variables and performed allelic turnover analyses with 22 geographical locations. The allelic turnover function predicts how frequencies of genetic variants in different geographical groups change with associated environmental variables. The overall allelic frequency turnover function for adaptive SNPs (cumulative importance of allele frequencies, y-axis) projected along these 5 factors (BIO4, BIO18, BIO5, BIO10 and BIO8, x-axis) was sigmoidal (Figure 6 B - F). Geographical group projections of cumulative importance of allele frequencies indicated distinct patterns between European and Japanese groups. The Japanese groups were in the ‘upper’ region of the sigmoid (higher cumulative importance of allele frequencies) while the European groups were in the ‘lower’ region of the sigmoid (lower cumulative importance of allele frequencies). The sigmoid of the allelic turnover function suggested that the Japanese groups are adapted to warmer temperatures and higher precipitation and European populations are adapted to cooler temperatures and lower precipitation (Figure S5). Furthermore, the European Atlantic populations are adapted to cooler wetter climate conditions than those in central Europe.

Analysis of sigmoid patterns indicates that cumulative allele frequency importance correlates with five climate variables (r=1, p<2.26e-16, Figure S6) across 22 geographical locations. To test if the longitude and or latitude influence adaptation, we projected cumulated importance of allelic frequencies (y axis) on to longitude and latitude of geographic groups (x axis). Projection of cumulative allele frequency importance onto longitude and latitude highlights associations with Maximum Temperature of Warmest Month (BIO5) and Mean Temperature of Warmest Quarter (BIO10), in European and Japanese groups (Figure S7A, B). In both groups, cumulative importance increases with decreasing latitude (North to South) for BIO5 and BIO10, while European groups also show an increase with increasing longitude (West to East) (Figure S7A B, Table S7). Conversely, Japanese groups show a slight increase with decreasing longitude (East to West) (Figure S7C, D, Table S7). The correlation extends to BIO4, reflecting seasonal climate variation, where cumulative importance increases with decreasing latitude in European groups (Figure S7E), and with increasing latitude and longitude in Japanese groups (Figure S7E, F). However, no positive correlation was found in Japan for BIO8 and BIO18 (Figure S7 G-J). We found that the cumulative importance tends to increase with increasing longitude for BIO8 (Figure S7J), while cumulative importance tends to increase with decreasing latitude for BIO8 and BIO18 in European groups. We conclude that the cumulated importance of allelic frequencies correlates with climate variables (BIO4, BIO5, and BIO10) that vary with latitude and longitude in Europe. Similarly, in Japan, the cumulated importance of allelic frequencies is correlated with climate variables (BIO4, BIO5, BIO10, BIO8, and BIO18) that vary with latitude and longitude. These data indicate that the frequencies of genetic variants in different geographic groups shift with different environmental variables.

### 7. Association mapping of the five most important climate variables

Genotype–environment association studies (GEAS) can discover genetic variants correlated with local climate factors [25]. We further applied GEAS to find association between genetic variants and climate variables. Before we performed GEAS, we calculated the heritability of 19 climate factors. The *H^2^* values exceeded 0.99 for BIO4, BIO5, BIO8 and BIO10, and was 0.89 for BIO18 (Table S8). Traits characterized by high heritability may be mapped by genetic association. These data therefore indicated that it should be possible to carry out GEAS analysis for BIO4, BIO5, BIO8, BIO10 and BIO18. GEAS was performed after employing a stringent Bonferroni-based threshold and Quantile-Quantile (Q-Q) plots method on the five most influential climate factors (Figure S8). We found six unlinked, significant SNPs associated with BIO4, ten unlinked significant SNPs associated with BIO5, four unlinked significant SNPs associated with BIO10 and nine unlinked significant SNPs associated with BIO8, respectively (Figure S8, Table S9). However, we did not find genetic variants significantly associated with BIO18 (Figure S8). These data demonstrate that specific genetic variants associated with putative roles in adaptation to local climate factors can be identified.

## DISCUSSION

We report considerable genetic diversity within local patches of *M. polymorpha* subspecies *ruderalis*, underscoring the high levels of variation present in some local populations. There are populations where every individual is genetically identical, and the population is entirely clonal derived from a single founder through asexual reproduction. Other populations are genetically diverse. The latter includes a mixture of different genotypes with some individuals being genetically identical. This demonstrates that both sexual and asexual reproduction is occurring in these populations. However, despite this substantial genetic diversity, we find no evidence of population structure across Europe. This lack of structure likely results from high levels of gene flow across the continent. Despite the lack of population structure on a continental scale across Europe, European and Japanese populations are genetically distinct. The genetic differentiation likely results from limited gene flow between European and Japanese populations.

The lack of population structure in Europe is likely due, in part at least, to long distance dispersal and gene flow throughout the continent. A similar lack of population structure has been identified in *Pinus sylvestris*[26], which is attributed to the production or large amounts of haploid pollen that can travel large distances, therefore mixing genetic diversity and such a panmixis could result in a lack of genetic differentiation on a continental scale. We speculate that, like the haploid pollen of *P. sylvestris* (30-100 µm diameter [27]), the potential for long-distance transport of the *M. polymorpha* haploid spores[7] may account for the lack of population structure in European *M. polymorpha* subspecies *ruderalis* populations. This lack of differentiation on a continental scale contrasts with what has been observed in the ruderal angiosperm, *A. thaliana*. The 1001 Genomes Consortium confirmed that the genetic distance between individuals reflects geographic distance[4]. Furthermore, the native Eurasian range of *A. thaliana* exhibits continuous isolation by distance at every geographic scale[2, 28]. We found no evidence that isolation by distance contributes to genetic distance in European *Marchantia polymorpha*. The difference in genetic structure between *M. polymorpha* and *A. thaliana* across Europe could be accounted by the different reproductive and dispersal strategies of these species: *M. polymorpha* is an outcrossing species and produces both very large number of small spores and fewer relatively heavy gemmae, while *A. thaliana* is largely inbreeding and produces fewer relatively heavy seed.

Mapping genetic diversity from plants growing in different environments allowed us to use genetic association for alleles that may underpin local adaptation. We report significant genetic adaptation to summer temperature and precipitation, particularly in the Japanese populations. Our demonstration that there is genetic basis to adaptation to climate factors using genome wide association, is consistent with previous findings in other plant species. For example, genome wide association studies in *A. thaliana* [29] and *Populus koreana* [30] identified genomic regions associated with adaptation to temperature and precipitation.

The discovery that summer temperature and precipitation play a significant role in shaping the genetic landscape of populations in *M. polymorpha* may reflect the requirement for water in the bryophyte life cycle. Bryophytes produce motile spermatozoids that swim through water to effect fertilization. It is therefore likely that the summer precipitation is a constraint on the completion of the *M. polymorpha life* sexual life cycle. Such a constraint would impose a considerable selection pressure that would be reflected in the presence of adaptive alleles that are correlated with temperature and precipitation during the time of year that *M. polymorpha* sexually reproduces. These alleles would be strongly selected for in ruderal environments which are generally open and relatively dry because of the drying effect of direct sunlight on the soil surface.

### Conclusions

We present a population genomic analysis of *M. polymorpha* natural selection and adaptation based on resequencing of unique 78 genomes of wild collections mainly from Europe and Japan. We find considerable genetic variation within patches that reproduced sexually and asexually. Other patches – derived from single individuals – reproduce exclusively through asexual reproduction are genetically monomorphic. Furthermore, we observed little differentiation among European populations while at an inter-continental level, Japanese and European populations are genetically distinct. This indicates that genetic diversity is distributed across Europe differently from other ruderal plants like *Arabidopsis thaliana.* Summer temperature and precipitation appear the most important climate factors affecting the frequency of adaptation alleles among these populations. Further sequencing of individuals in populations from all continents and isolated islands will allow higher resolution mapping of alleles associated with local adaptation.

## MATERIALS AND METHODS

### Sample collection and sterilization

*Marchantia polymorpha* subsp*. ruderalis* accessions were collected from 10 European countries, Japan, US and Canada from August 2021 to June 2023 (Table S1). Collected tissue samples were precleaned individually with tap water then grown on soil (2:1 ratio of Neuhaus Huminsubstrat N3 from Klasmann-Deilmann GmbH and fine vermiculite) in Sac O2 Microboxes (OV80+OVD80 #40 NG/NP) under standard growth conditions (23°C, continuous white light 50 – 60 µmol m-2 s-1). The plants were allowed to grow until they developed gemmae. Gemmae were surface sterilized in 0.1 - 1% sodium hypochlorite containing 0.01% Triton-X for 5 minutes, washed in sterile dH20 for 1 minute and finally rinsed three times in sterile dH20. Gemmae were transferred to sterile plates containing solid 0.5x Gamborg medium (1.5 g/L B5 Gamborg, 0.5 g/L MES hydrate, 1% plant agar, pH adjusted to 5.5) supplemented with Cefotaxime (100mg/l) and grown under standard growth conditions.

Thallus sterilization was performed for *M. polymorpha* subsp. ruderalis samples which didn’t develop gemmae cups. Thallus explants of 0.4 - 0.8 cm² were sterilized in 1% NaDCC (Sodium dichloroisocyanurate) with 0.01% Triton-X for 1 minute, washed in sterile dH20 for another minute and rinsed three times with sterile dH2O before drying on filter paper. Following this surface sterilization process the thallus explants were moved to solid 0.5x Gamborg medium with Cefotaxime (100mg/l) and grown under standard growth conditions. For samples which didn’t remain free of visible contamination, the thallus sterilization procedure was repeated after cleaning the explants in 70% EtOH for about 5 seconds.

### Genomic DNA sequencing

Axenic natural accession lines were grown from gemmae on solid 0.5x Gamborg medium under standard growth conditions. After three weeks of growth, 70-80 mg of plant tissue was harvested and flash frozen in liquid nitrogen. Genomic DNA was extracted using the Qiagen DNeasy Plant Pro Kit (Cat. # 69204) according to the kit protocol with an additional incubation (65°C for 10min) prior to the bead-beating step (using Retsch mill at 30 Hz, 4min) to improve the yield. The gDNA was eluted in EB buffer.

The gDNA concentration was measured using a Qubit 4.0 fluorometer according to the instruction manuals. Genomic DNA samples were sent for sequencing to the Next Generation Sequencing Facility at Vienna BioCenter Core Facilities (VBCF), member of the Vienna BioCenter (VBC), Austria. DNA Libraries were prepared using NEBNext® Ultra™ II FS DNA Library Prep Kit for Illumina and fragment size was determined using a BioLabTech Fragment Analyzer. The DNA was sequenced on Illumina NovaSeq SP / NovaSeq S1 / NovaSeq S4 flowcells using 150 bp paired-end reads.

### Read mapping, variant calling, and clones removing

Adapters and low-quality reads (q < 20) were removed using Trimmomatic (version 0.39)[31]. Trimmed reads with a length shorter than 36 bp were discarded. The remaining clean reads were then aligned to the Tak1 V6.1 genome using BWA-MEM (version 0.7.17-r1188) with default parameters [32]. Alignment quality was assessed using SAMtools (MAPQ = 30) [33] and RSeQC (version 2.6.4)[34].

For variant calling, the Genome Analysis Toolkit Haplotype Caller (version 4.2.6.1) [35] was employed. Initially, per-sample GVCF files were generated, followed by joint genotyping with GenotypeGVCFs (using -ploidy 1). A stringent filtering approach was applied based on the removal of tails in the variant distributions, as suggested by a previous study[17]. According to this approach, SNPs with the following criteria were excluded: ’QUAL < 43.4 || DP < 68 || DP > 1295 || MQ < 37.4 || SOR > 3.61 || QD < 4.81 || FS > 13.2 || MQRankSum < -3.72 || ReadPosRankSum < -1.96 || ReadPosRankSum > 2.23’. Subsequently, VCFtools [36] was utilized to obtain pure SNPs with the following parameters: --min-alleles 2 --max-alleles 2 --hwe 1e-06 -- maf 0.02 --max-missing 0.9.

### Removal of clonal accessions

Hamming distance, calculated by Plink [37], was used to calculate the genome distance between individuals based on clean SNPs. We then computed the hamming distance among the 209 accessions using all autosomal SNPs. The genome distance (GD) distribution revealed two distinct peaks (Figure S1B): peak A corresponding to a GD of less than 0.01, and peak B corresponding to a GD of 0.24. We also plotted the GD distribution within each population and observed a similar pattern (Figure 1B). Therefore, we concluded that pairs of accessions with GD values less than 0.01 (peak A) were considered to be clones. Following the removal of these clones, a total of 78 accessions were retained for downstream analysis.

Given that *M. polymorpha* vegetatively reproduces forming clones, we removed clones from our analyses because they would skew, erroneously the conclusions. Our analysis was carried out using genetically unique individuals by removing clones. The analysis revealed the existence of two genetically distinct groups: European and Japanese populations. However, if all individuals 209 –including unique and clonal individuals – were included in the population structure analysis, the results changed. The PCA analysis revealed four groups (Figure S9 A ): Japanese, most European accessions, Sopron (Hungary), and MaG (Austria). However, we know that the Sopron (24 individuals) and MaG (24 individuals) populations are clonal; pairwise genome distance within Sopron and MaG populations was less than 0.01. Therefore, the inclusion of clonal individuals would have biased our analysis. We also found that the large number individuals of clonal populations of Sopron and MaG also affect the admixture result. We observed that the Sopron and MaG populations retained their distinct ancestries from K=2 to K=9 (Figure S9 B, Figure S9 C). Previous research highlighted that clonal populations could cluster tightly, skewing the principal components and obscuring meaningful patterns of genetic variation in non-clonal populations[38]. Inclusion of clones in the analysis can therefore misrepresent the true ancestral proportions of the broader population[38]. Thus, failing to remove clonal accessions introduces bias in population structure analysis. This was evident when the population structure clearly changed after removing all clonal populations. For instance, in the non-clonal population structure of 78 accessions, the Sopron and MaG accessions were classified within the European group (Figure S9D). Therefore, it is crucial to remove clonal accessions with pair wise genome distance less than 0.01 before conducting population genomics studies, especially in haploid-dominant life cycle plants.

### Population structure, genetic diversity and differentiation analyses

Population structure analyses were performed with two methods, PCA and admixture. PCA analysis was performed on the pure SNP set using GCTA (v1.93.2 beta)[39]. ADMIXTURE (v 1.3)[40] with cross-validation was used to investigate population structure across all individuals, with the number of clusters (K) being set from 1 to 9. The best K is 2 based on the smallest CV value.

Nucleotide diversity Pi), pairwise nucleotide diversity (Dxy) and genetic differentiation (Fst) were calculated using pixy [41]with a nonoverlapping 20-kb sliding window across the genome suggested by previous study on haploid bryophyte and fungus [42, 43]. The input to pixy was an all-sites VCF containing both variant and non-variant sites generated by GATK (‘GenotypeGVCFs -all-sites’) suggested by Pixy manual. The median values across all sliding windows were reported for Pi, Dxy, and Fst based on previous study[17, 30].

### Isolation by distance (IBD) and Isolation by environment (IBE) analysis

IBD was analyzed based on relationship between genetic distance and geographical distance. Genetic distance was defined with Fst/(1-Fst ) based on previous study[28, 30, 44]. Geographical distance was calculated using pair-wise Latitude and Longitude information with Geosphere (https://github.com/rspatial/geosphere). The correlation between genetic distance and geographical distance was tested using spearman test. IBE was analyzed based on relationship between genetic distance and environment distance. Environment distance was evaluated by euclidean distance using 19 climate factors downloaded from Worldclim (2.1)[22]. The correlation between genetic distance and environment distance was tested using spearman test.

### Linkage disequilibrium and demographic history analysis

Linkage disequilibrium (LD) was assessed using PopLDdecay (version 3.42; https://github.com/BGI-shenzhen/PopLDdecay) with default parameters based on previous study[19, 20]. The LD decay was visualized by Plot_MutiPop.pl function implemented in PopLDdecay. LD was quantified as the pairwise SNP distance at which the allele frequency correlation (r2) dropped to half of its maximum value.

To elucidate the historical changes in the effective population size of Marchantia populations, demographic analyses were performed employing SMC++ (version 1.15.5)[45]. The effective population size denotes the number of breeding individuals in an idealized population that would result in the observed genetic diversity. For each population, SNPs were called using VCFtools (--max-missing 1), and subsequently, smc++ vcf2smc was executed for each autosome. Estimated population sizes were computed using smc++ estimate, incorporating a mutation rate of 2.5e-9 mutations per site per generation[46]. The effective population size per generation was scaled based on an estimated generation time of one generation per year.

### Environment-associated genetic variants identification, heritability estimation, and gradient forest analysis

Nineteen climate variables were retrieved from the WorldClim v2.1 database [22]), encompassing BIO1 (Annual Mean Temperature), BIO2 (Mean Diurnal Range), BIO3 (Isothermality), BIO4 (Temperature Seasonality), BIO5 (Max Temperature of Warmest Month), BIO6 (Min Temperature of Coldest Month), BIO7 (Temperature Annual Range), BIO8 (Mean Temperature of Wettest Quarter), BIO9 (Mean Temperature of Driest Quarter), BIO10 (Mean Temperature of Warmest Quarter), BIO11 (Mean Temperature of Coldest Quarter), BIO12 (Annual Precipitation), BIO13 (Precipitation of Wettest Month), BIO14 (Precipitation of Driest Month), BIO15 (Precipitation Seasonality), BIO16 (Precipitation of Wettest Quarter), BIO17 (Precipitation of Driest Quarter), BIO18 (Precipitation of Warmest Quarter), BIO19 (Precipitation of Coldest Quarter). Climate-associated SNPs were identified using the LFMM method, implementing a Latent Factor Mixed Model. Considering the number of ancestry clusters inferred with ADMIXTURE, LFMM was executed with two latent factors to address population structure in the genotype data. Adaptive SNPs were selected based on the top 0.0005% adjusted P value of all SNPs from LFMM, resulting in 2191 adaptive SNPs across the 19 climate variables. Significant SNPs were identified using a Bonferroni-based threshold and the Quantile-Quantile (Q-Q) method.

Heritability estimation are based on clean SNPs by GCTA-GREML [39]. To be briefly, 1) Genomic Relationship Matrix construction: GCTA starts by creating a genomic relationship matrix (GRM), which quantifies the genetic similarity between pairs of individuals based on genome-wide SNP data. This matrix represents the proportion of shared genetic material between individuals. 2) Phenotypic Variance Decomposition: Using the GRM and the bioclimate factors, GCTA applies the REML algorithm to partition the total phenotypic variance (*V_p_*) into two components: Genetic variance (*V_g_*): The proportion of variance due to additive genetic factors. Environmental variance (*V_e_*): The proportion of variance due to non-genetic factors. 3) Heritability Estimation: The narrow-sense heritability (*h^2^*) is calculated as the ratio of genetic variance to total phenotypic variance (*V_g_/V_p_*).

Gradient forest (GF), a nonparametric, machine-learning regression tree method, utilized SNP allele frequencies and climatic data to identify environmental gradients associated with genetic variation and determine how allele frequencies change along those gradients [23]. Allelic frequencies for 2191 SNPs were calculated using VCFtools across 22 geographical groups. Weighted importance was assessed based on the 19 Bioclimate variables and allelic frequencies using GF. Cumulative importance of allelic frequency was calculated for the five most significant climate variables, projecting onto the geographical groups. This cumulative importance analysis is followed the protocol based on previous study[47].

## RESOURCE AVAILABILITY

Lead contact Further information and requests for resources and reagents should be directed to and will be fulfilled by the lead contact, Liam Dolan.

### Data and code availability

High through-put sequencing data generated in this study have been deposited at NCBI SRA with code PRJNA1127023 and are publicly available as of the date of publication (reviewer link: https://dataview.ncbi.nlm.nih.gov/object/PRJNA1127023?reviewer=hctoksdj83ba550ts1unjnj0er). All code used in this study is available on GitHub (https://github.com/wushyer/Population-genomics-of-Marchantia-polymorpha-sub.-ruderalis).

## ACKNOWLEDGMENTS

We thank Lab Support of GMI/IMBA/IMP and the VBCF Plant Sciences unit for their support and the Next Generation Sequencing Facility for generating the DNA sequencing data (VBCF). We thank Jonathan Terhorst suggestions of using SMC++ on *M.polymorpha* sub*. ruderalis* population history analysis. We thank Masanobu Yoshidomi and Takahiro Mochizuki for help in collecting plant materials in Japan. This work was funded by a grant from the Austrian Academy of Sciences to the Gregor Mendel Institute and a European Research Council advanced grant DENOVO-P (project no. 787613) to L.D, and Marie Skłodowska-Curie Grant Agreement no. 847548 (VIP2) to S.W.

## AUTHOR CONTRIBUTIONS

S.W. and L.D. designed the project. S.W. carried out the project with help from K.S. K.J. and S.A. performed all the accessions cultivation, which were collected by J.R., M.S., T.H., and F.B. K.J. performed genome sequencing and S.W. analyzed the data. S.W. and L.D. wrote the manuscript draft. The draft was revised with input from F.B. and K.S.

## SUPPLEMENTARY MATERIAL

Figure S1 Genome distances distribution among 12 populations accessions, with the dashed line indicating the threshold for identifying clones.

Figure S2 Genetic profile of the 78 accessions. A) Admixture result from K1 to K9. B) K number distribution of two selected populations.

Figure S3 Pi and Fst distribution in genome and autosome, respectively. A) Nucleotide diversity among Europe and Asia populations was analyzed on a genome-wide scale. Nucleotide diversity was measured in 20 kb windows. B) The distribution of Pi values for each autosome is shown in boxplot with chr1 to chr8. The solid horizontal lines indicate the median pi values. C) Genome-wide genetic differentiation between Asia and Europe populations was compared using pairwise Fst measurements in 20 kb windows. D) Density plots of the distribution of Fst values for each autosome, with chr1 to chr8 arranged from bottom to top. The solid vertical lines represent the median FST values. The dashed line is the Fst of whole genome.

Figure S4 Spearman’s correlation coefficient (two-sided test) of 19 climate factors.

Figure S5 Phenotype distribution of five most significant bioclimate factors including BIO4, BIO5, BIO10, and BIO8.

Figure S6 Correlation between Cumulated importance of allele frequencies and 5 most significant bioclimate factors.

Figure S7 Correlation between cumulated importance of allele frequencies and latitude/longitude in 5 most significant bioclimate factors. A-B) Correlation between cumulated importance of allele frequencies and latitude/longitude in BIO10. C-D) Correlation between cumulated importance of allele frequencies and latitude/longitude in BIO5. E-F) Correlation between cumulated importance of allele frequencies and latitude/longitude in BIO4. G-H) Correlation between cumulated importance of allele frequencies and latitude/longitude in BIO18. I-J) Correlation between cumulated importance of allele frequencies and latitude/longitude in BIO8.

Figure S8 QQ plots and GEAS results for BIO4, BIO5, BIO10, and BIO8. The QQ plot visually represents the deviation of observed P-values from the null hypothesis. For each SNP, observed P-values are sorted from largest to smallest and plotted against expected values derived from a theoretical χ²-distribution. Agreement between observed and expected values is indicated by points clustering around the middle line between the x-axis and y-axis. In the GEAS results, the dashed line signifies the cutoff for significant variants. Panels C and D show QQ plots and GEAS results for BIO5. Panels E and F show QQ plots and GEAS results for BIO10. Panels G and H show QQ plots and GEAS results for BIO8. Panels I and J show QQ plots and GEAS results for BIO18.

Figure S9 Population structure of 209 accessions including clones. A) Principal component analysis (PCA) of 209 accessions revealed four distinct groups: Japanese, most European accessions, Sopron (Hungary), and MaG (Austria). B)

ACross-validation (CV) error analysis of the admixture results showed no optimal number of groups. C) Genetic profiles of the 209 accessions from K2 to K9. Two clonal populations maintained their genetic profiles from K2 to K9, highlighting the distinct characteristics of clonal populations. D) PCA of 78 non-clonal accessions, revealing a different population structure compared to the analysis that included clonal accessions.

Table S1 General information of 209 accessions.

Table S2 Quality control of 209 accessions.

Table S3 Pairwise genome distance of 209 accessions.

Table S4 78 nonclone accessions.

Table S5 Pi value of 7 European populations.

Table S6 Median Fst values across different genomic regions of autosomes

Table S7 Geographical location of the dot in BIO4, BIO5 and BIO10 of Figure 8.

Table S8 SNP heritability with 19 climate factors

Table S9 Significant variant of GEAS in BIO4, BIO5, BIO10 and BIO8.

